# SAVER: Gene expression recovery for UMI-based single cell RNA sequencing

**DOI:** 10.1101/138677

**Authors:** Mo Huang, Jingshu Wang, Eduardo Torre, Hannah Dueck, Sydney Shaffer, Roberto Bonasio, John Murray, Arjun Raj, Mingyao Li, Nancy R. Zhang

**Affiliations:** Department of Statistics, University of Pennsylvania, Philadelphia, PA; Department of Bioengineering, University of Pennsylvania, Philadelphia, PA; Department of Genetics, University of Pennyslvania, Philadelphia, PA; Department of Cell and Developmental Biology, University of Pennsylvania, Philadelphia, PA; Department of Biostatistics and Epidemiology, University of Pennsylvania, Philadelphia, PA

## Abstract

Rapid advances in massively parallel single cell RNA sequencing (scRNA-seq) is paving the way for high-resolution single cell profiling of biological samples. In most scRNA-seq studies, only a small fraction of the transcripts present in each cell are sequenced. The efficiency, that is, the proportion of transcripts in the cell that are sequenced, can be especially low in highly parallelized experiments where the number of reads allocated for each cell is small. This leads to unreliable quantification of lowly and moderately expressed genes, resulting in extremely sparse data and hindering downstream analysis. To address this challenge, we introduce SAVER (Single-cell Analysis Via Expression Recovery), an expression recovery method for scRNA-seq that borrows information across genes and cells to impute the zeros as well as to improve the expression estimates for all genes. We show, by comparison to RNA fluorescence in situ hybridization (FISH) and by data down-sampling experiments, that SAVER reliably recovers cell-specific gene expression concentrations, cross-cell gene expression distributions, and gene-to-gene and cell-to-cell correlations. This improves the power and accuracy of any downstream analysis involving genes with low to moderate expression.

## Introduction

A primary challenge in the analysis of scRNA-seq data is the low capturing and sequencing efficiency affecting each cell, which leads to a large proportion of genes, often exceeding 90%, with zero or low read count. Although many of the observed zero counts reflect true zero expression, a considerable fraction is due to technical factors such as capture and sequencing efficiency. The overall efficiency of current scRNA-seq protocols can vary between <1% to >60% across cells, depending on the method used^1^.

Existing studies have adopted varying approaches to mitigate the noise caused by low efficiency. In differential expression and cell type classification, transcripts expressed in a cell but not detected due to technical limitations, also known as dropouts, are sometimes accounted for by a zero-inflated model^2–4^. Recently, methods such as MAGIC^5^ and scImpute^6^ have been developed to directly estimate the true expression levels. Both MAGIC and scImpute rely on pooling the data for each gene across similar cells. However, as we show later in results, this can lead to over-smoothing and may remove natural cell-to-cell stochasticity in gene expression, which has been shown to lead to biologically meaningful variations in gene expression, even across cells of the same type^7,8^ or of the same cell line^9,10^. In addition, MAGIC and scImpute do not provide a measure of uncertainty for their estimated values.

Here, we propose SAVER (Single-cell Analysis Via Expression Recovery), a method that takes advantage of gene-to-gene relationships to recover the true expression level of each gene in each cell, removing technical variation while retaining biological variation across cells (https://github.com/mohuangx/SAVER). SAVER receives as input a post-QC scRNA-seq dataset with unique molecule index (UMI) counts, see Methods for the recommended data pre-processing. SAVER assumes that the count of each gene in each cell follows a Poisson-Gamma mixture, also known as a negative binomial model. Instead of specifying the Gamma prior, we estimate the prior parameters in an empirical Bayes-like approach with a Poisson Lasso regression^11^ using the expression of other genes as predictors. Once the prior parameters are estimated, SAVER outputs the posterior distribution of the true expression, which quantifies estimation uncertainty, and the posterior mean is used as the SAVER recovered expression value (Methods).

We first evaluate the performance of SAVER through comparisons of Drop-seq and RNA FISH on a melanoma cell line. We demonstrate through down-sampling experiments that the observed expression is distorted by efficiency loss, but that the true expression profiles can be recovered using SAVER. Finally, we apply SAVER to a mouse cortex dataset to show that SAVER can identify true cell types with only a fraction of the cells.

## Results

### Overview of SAVER model and interpretation

SAVER is based on adaptive shrinkage to a multi-gene prediction model (Fig. 1a). Let *Y*_*cg*_ denote the observed UMI count of gene *g* on cell *c*. We model *Y*_*cg*_as

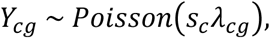

**Figure 1.**
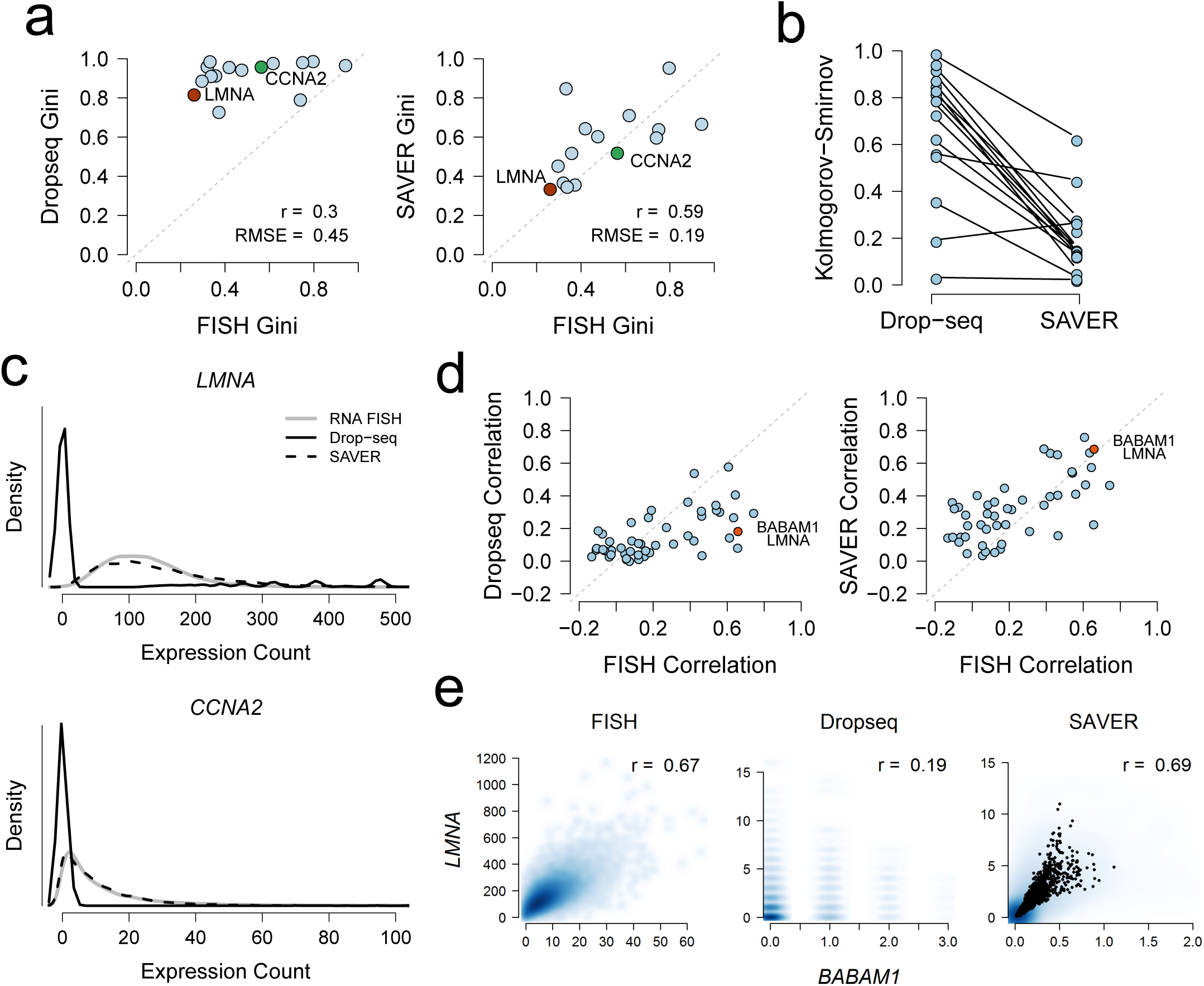
RNA FISH validation of SAVER results on Drop-seq data for 15 genes. (**a**) Comparison of Gini coefficient for each gene between FISH and Drop-seq (left) and between FISH and SAVER recovered values (right). (**b**) Comparison of SAVER and Drop-seq Kolmogorov-Smirnov distance to FISH distributions for the 15 genes. (**c**) Kernel density estimates of cross-cell expression distribution of *LMNA* (upper) and *CCNA2* (lower). (**d**) Comparison of pair-wise gene correlations computed from Drop-seq original counts (left) and from SAVER recovered values (right) with those computed from FISH counts. SAVER is able to recover the depressed gene correlations in Drop-seq. (**e**) Scatterplots of expression levels between *BABAM1* and *LMNA*.

where *λ*_*cg*_ is the true expression level of gene *g* in cell *c*, and *s*_*c*_ is a cell-specific size factor which will be described below. The Poisson distribution after controlling for biological variation between cells has been shown previously by bulk-RNA splitting experiments to be a reasonable approximation for observed gene counts. Our goal is to recover *λ*_*cg*_ with the help of a prediction *μ*_*cg*_ based on the observed expression of a set of informative genes *S*_*g*_ in the same cell (Methods). The accuracy of *μ*_*cg*_ in predicting *λ*_*cg*_ differs across genes — genes that play central roles in pathways are easier to predict, whereas genes that are not coordinated with other genes are harder to predict. To account for prediction uncertainty, we assume for *λ*_*cg*_ a gamma prior with mean set to the prediction *μ*_*cg*_ and with dispersion parameter *ϕ*_*g*_. The dispersion parameter quantifies how well the expression level of gene *g*is predicted by *μ*cg. After maximum-likelihood estimation of *ϕ*_*g*_ and reparameterization, let 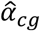 and 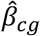 be the estimated shape and rate parameters, respectively, for the prior gamma distribution. Then, the posterior distribution of *λ*_*cg*_ is also gamma distributed with shape parameter 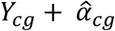 and rate parameter 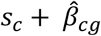. The SAVER recovered gene expression is the posterior mean,

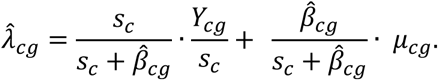

As seen from the above equation, the recovered expression 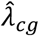 is a weighted average of the normalized observed counts *Y*_*cg*_ */s*_*c*_ and the prediction *μ*_*cg*_. The weights are a function of the size factor *s*_*c*_ and, through the 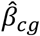 term, the gene’s predictability 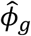 and its prediction *μ*_*cg*_. Genes for which the prediction is more trustworthy (small 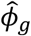) have larger weight on the prediction *μ*_*cg*_. Genes with higher expression have larger weight on the observed counts and rely less on the prediction. Cells with higher coverage have more reliable observed counts and also rely less on the prediction. Figure 1b shows example scenarios.

Interpretation of *λ*_*cg*_ depends on how the size factor *s*_*c*_ is defined and computed. There are two scenarios. In what is perhaps the simpler scenario, assume that the efficiency loss, that is, the proportion of original transcripts that are sequenced and observed, is known or can be estimated through external spike-ins. If *s*_*c*_ were defined as the cell-specific efficiency loss, then *λ*_*cg*_ would represent the absolute count of gene *g*in cell *c*. The second scenario assumes that the efficiency loss is not known, in which case *s*_*c*_ can be set to a normalization factor such as library size or scRNA-seq normalization factors^12,13^. In this case, *λ*_*cg*_ represents a gene concentration or relative expression. Which scenario applies depends on the objective of the study and the availability and quality of spike-ins.

It is important to note that SAVER outputs the posterior distribution for *λ*_*cg*_, not just its posterior mean. The posterior distribution quantifies the uncertainty in our estimate of *λ*_*cg*_, and it is crucial to incorporate this uncertainty in downstream analyses. We demonstrate the use of this posterior distribution in two ways. First, to recover the cross-cell expression distribution of a given gene, we sample from the posterior of *λ*_*cg*_ for each cell instead of simply using the posterior mean 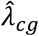. Second, in estimating gene-to-gene correlations, the sample correlation of the recovered estimates 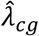 tends to overestimate the true values. We developed an adjusted measure of correlation which takes into account the uncertainty in *λ*_*cg*_ (see Methods).

### Using RNA FISH as gold standard, SAVER recovers gene expression distributions

First, we assessed SAVER’s accuracy by comparing the distribution of SAVER estimates to distributions obtained by RNA FISH in data from Torre and Dueck et al.^14^ In this study, Drop-seq^15^ was used to sequence 8,498 cells from a melanoma cell line. In addition, RNA FISH measurements of 26 drug resistance markers and housekeeping genes were obtained across 7,000 to 88,000 cells from the same cell line. After filtering, 15 genes overlapped between the Drop-seq and FISH datasets; these overlapping genes are representative of all genes in mean and % non-zero expression, but have slightly higher dispersion (Supplementary Fig. 1). Since FISH and scRNA-seq were performed on different cells, the FISH and scRNA-seq derived estimates can only be compared in distribution. We applied SAVER to the Drop-seq data and calculated the Gini coefficient^16^, a measure of gene expression variability, for the FISH, Drop-seq, and SAVER results for these 15 overlapping genes. The Gini coefficient has been shown to be a useful measure for identifying rare cell types and sporadically expressed genes in the original FISH-based study of this cell line^10^, and thus, accurate recovery of the Gini coefficient would allow the same analysis to be performed with scRNA-seq. For all genes, SAVER effectively recovered the FISH Gini coefficient, which Drop-seq grossly overestimates (Fig. 1a).

To evaluate the similarity of the distributions, we also calculated the Kolmogorov-Smirnov (KS) distance between FISH and Drop-seq and between FISH and SAVER for each gene (Fig. 1b). The distributions of the SAVER recovered expression values match much more closely with the FISH distributions than the distributions of the Drop-seq counts, as demonstrated by the reduced KS distances and by direct overlays of density plots for each gene (Fig. 1c, Supplementary Fig. 2). In comparison, Gini estimates and recovered distribution obtained from MAGIC match poorly with the FISH estimates; scImpute performs better than MAGIC at recovering the Gini coefficient and distributions but not as well as SAVER (Supplementary Fig. 3a-c). Accurate recovery of gene expression distribution is important for identifying rare cell types, identifying highly variable genes, and studying transcriptional bursting.

**Figure 2.**
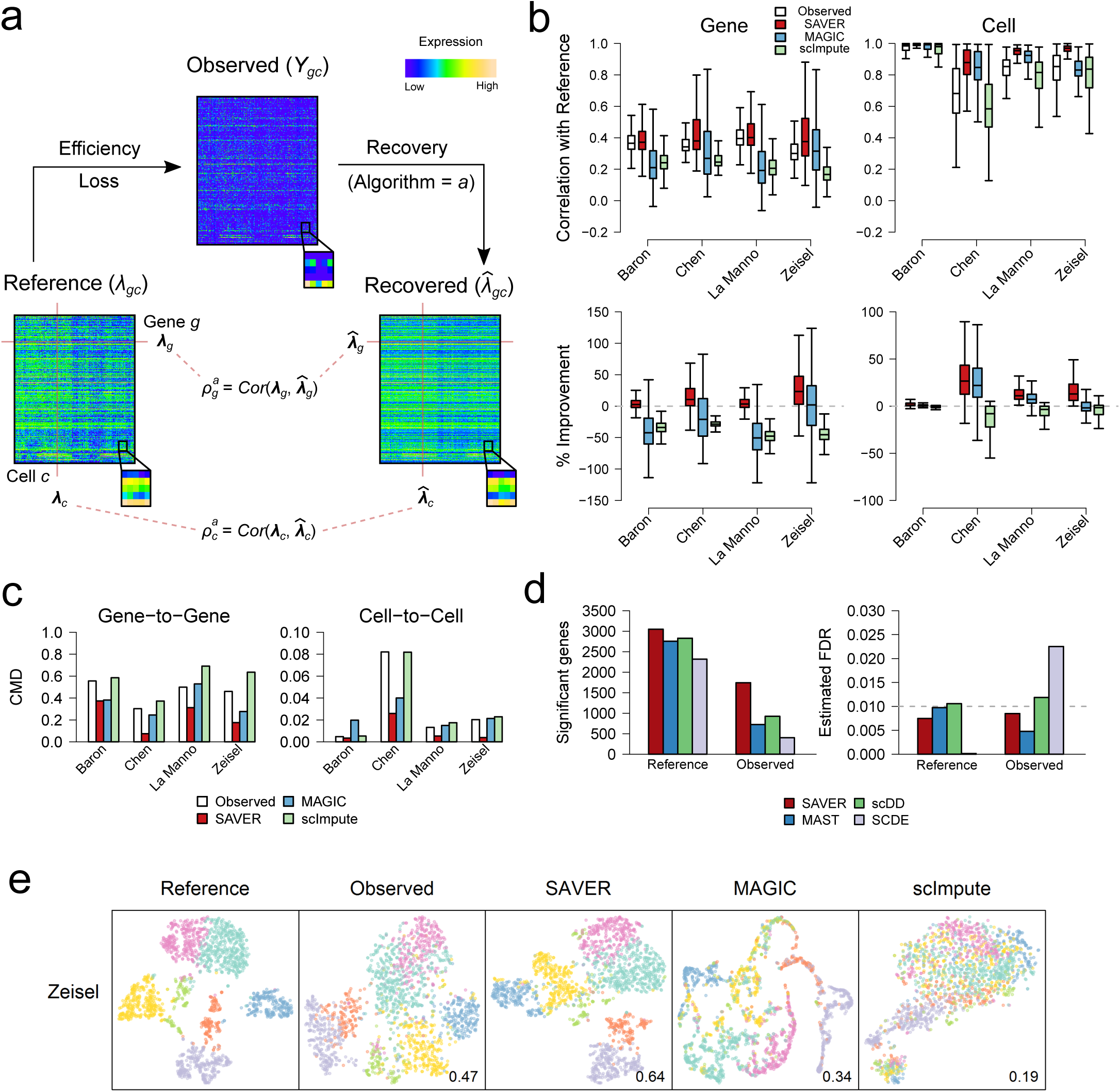
Evaluation of SAVER by down-sampled mouse brain dataset. (**a**) Schematic of down-sampling experiment. (**b**) Performance of algorithms measured by correlation with reference, on the gene level (left) and on the cell level (right). Percentage improvement over using the observed data is shown in the lower two panels. SAVER is more closely correlated with the truth across both genes and cells. (**c**) Comparison of gene-to-gene (left) and cell-to-cell (right) correlation matrices of recovered values with the true correlation matrices, as measured by correlation matrix distance (CMD). (**d**) Differential expression (DE) analysis between CA1Pyr1 cells (*n* = 351) and CA1Py2 cells (*n* = 389). SAVER yields more significant genes in the down-sampled data (left), while still controlling false discovery rate at 0.01 (right). (**e**) Cell clustering and t-SNE visualization of the Zeisel dataset. Jaccard index of the down-sampled observed dataset and recovery methods as compared to the reference classification is shown.

**Figure 3.**
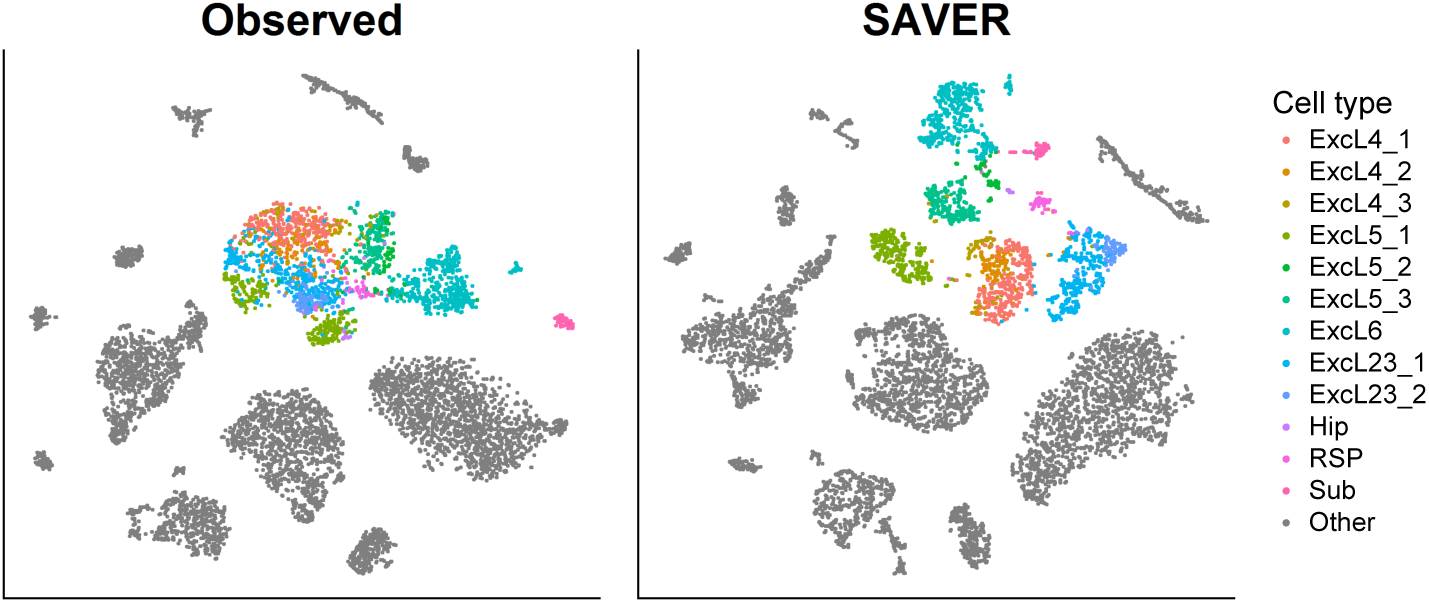
t-SNE visualization of 7,387 Hrvatin mouse visual cortex data for the observed data (left) and SAVER (right). Colored cells are excitatory neurons with each color representing subtypes determined by Hrvatin et al. using 47,209 cells. SAVER accurately separated the subtypes using a fraction of the original cells.

### SAVER recovers gene-to-gene relationships that are validated by RNA FISH

Not only is SAVER capable of recovering gene expression distributions and distribution-level features, it is also able to recover true biological gene-to-gene correlations that are observed in FISH but dampened in Drop-seq (Fig. 1d). For example, SAVER can recover the strong correlation between housekeeping genes *BABAM1* and *LMNA*, which is lost in the Drop-seq data (Fig. 1e). In comparison, the correlations derived from MAGIC results are much higher than those derived from FISH, suggesting that MAGIC induces spurious correlation. On the other hand, scImpute averages the correlations, leading to biased estimates of the true correlation (Supplementary Fig. 3d-e). The fact that SAVER does not introduce spurious correlation for gene pairs that have no biological correlation is further demonstrated by a permutation study (Supplementary Note 4), which shows that for such gene pairs the correlation estimates are shrunk to zero by SAVER, but inflated by MAGIC and scImpute (Supplementary Fig. 4). SAVER’s ability to estimate gene-pair correlations without over-smoothing allows for more accurate gene interaction and network analysis.

### SAVER recovers cell-specific gene expression values

Next, we evaluated whether SAVER can accurately recover the true expression level within each individual cell for each gene, in addition to recovery of distributions and correlations. Since it is difficult to determine the actual number of mRNA molecules in each cell prior to isolation and capture in an scRNA-seq experiment, to generate realistic benchmarking datasets, we performed down-sampling experiments on four datasets^17–20^. For each dataset, we first selected a subset of highly expressed genes and cells to act as the reference dataset, which we treat as the true expression. We then simulated the capture and sequencing process according to Poisson sampling at low efficiencies while introducing cell-to-cell variability in library size (Fig. 2a, Methods). The observed down-sampled datasets were created to have approximately the same percentage of non-zero expression as the original, full dataset (Supplementary Table 1). We ran SAVER, MAGIC, and scImpute on each of the observed datasets, as well as conventional missing data imputation algorithms (K nearest neighbors^21^, singular value decomposition^22^, random forest^23^).

To evaluate the performance of each method, we calculated the Pearson gene-wise correlation 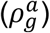 across cells and the cell-wise correlation 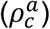 across genes between the reference and observed data, as well as between the reference and recovered datasets (Fig. 2a,b). The gene-wise correlations between the observed data and the reference data mostly range from 0.2 to 0.6 across all datasets. There is more variability in the cell-wise correlations across datasets, but these are generally higher than the gene-wise correlations. SAVER improves on both the gene-wise and cell-wise correlations across all datasets, while MAGIC, scImpute, and conventional missing data imputation algorithms usually perform worse than using the observed data (Fig. 2b, Supplementary Fig. 5,6a).

Next, we assessed the recovery of gene-to-gene and cell-to-cell correlation matrices, necessary for gene network reconstruction and cell type identification respectively. To compare, we calculated the correlation matrix distance (CMD)^24^ between the reference matrix and the observed/recovered matrix (Fig. 2c). SAVER lowers the gene-to-gene and cell-to-cell CMD for all datasets, MAGIC and scImpute perform similarly as the observed, and conventional missing data imputation algorithms perform worse than observed (Supplementary Fig. 6b).

### SAVER improves differential expression analysis and cell clustering

To investigate the effect of SAVER on downstream analyses, we performed differential expression and cell clustering on the down-sampled data. In the Zeisel study, two subclasses of cells — 351 CAPyr1 and 389 CA1Pyr2 cells — were identified by BackSPIN, a biclustering algorithm. We performed differential expression analysis of these two subclasses using several differential expression methods^2,3,25^. After down-sampling, the number of differentially expressed genes detected is much lower than for the reference, but SAVER is able to detect the most genes in the down-sampled data set while maintaining accurate FDR control (Fig. 2d, Supplementary Table 2).

Next, we performed cell clustering on the reference, observed, and recovered datasets using Seurat^26^. The reference-derived cell type clusters were treated as the truth and clustering accuracy on the observed and recovered datasets was assessed by the Jaccard index^27^ and by t-SNE^28^ visualization. SAVER achieves a higher Jaccard index than the observed for all datasets, while MAGIC and scImpute have a consistently lower Jaccard index (Fig. 2e, Supplementary Fig. 7). Even though the Jaccard index for SAVER in the Chen and La Manno datasets are only slightly higher than the observed, the t-SNE plots reveal that SAVER clustering of the cells is a more accurate representation of the reference data than the observed. SAVER also gives more stable results across different numbers of principal components, a critical parameter choice for dimension reduction in Seurat prior to the application of t-SNE (Supplementary Fig. 8).

### Illustration of SAVER in cell type identification of mouse cortex cells

Finally, we demonstrated SAVER in the analysis of a mouse visual cortex dataset where 47,209 cells were classified into main cell types and subtypes through extensive analysis^29^. We applied SAVER to a random subset of 7,387 cells (see Methods) and performed t-SNE visualization of the observed versus the SAVER-recovered cells (Fig. 3). A population of excitatory neurons is highlighted, and the individual subtypes are colored according to labels given by Hrvatin et al. In the t-SNE plot of the original counts, the subtypes are not well separated and are mostly indistinguishable. SAVER is able to distinguish the individual subtypes with clear separation. This example is common in our general experience with SAVER: It does not affect well-separated cell types, but identifies cell types and states for which the evidence in the original data may be weak.

## Discussion

We have described SAVER, an expression recovery method for scRNA-seq. SAVER aims to recover true gene expression patterns by removing technical variation while retaining biological variation. SAVER uses the observed gene counts to form a prediction model for each gene and then uses a weighted average of the observed count and the prediction to estimate the true expression of the gene. The weights balance our confidence in the prediction with our confidence in the observed counts. In addition, SAVER provides a posterior distribution which reflects the uncertainty in the SAVER estimate and which can be sampled from for distributional analysis.

We have shown that SAVER is able to accurately recover both population-level expression distributions and cell-level gene expression values, both of which are necessary for effective downstream analyses. Additional in-depth exploration in Supplementary Note 3 reveals how the performance of SAVER depends on factors such as sequencing depth, number of cells, and cell composition. In almost all scenarios, analyses using SAVER estimates improves upon analyses using the original counts, while in the worst case scenario, SAVER does not seem to hurt. The robust performance of SAVER is due to its adaptive estimation of gene-level dispersion parameters and its cross-validation-based model selection, which safeguard against unnecessary model complexity. By reducing noise and amplifying true biological relationships, SAVER improves the signal for downstream analyses.

## Acknowledgments

M.H. and N.R.Z. acknowledge support from NIH R01HG006137 and R01GM125301. M.H. is also supported by the National Science Foundation Graduate Research Fellowship Program under Grant No. DGE-1321851. Any opinions, findings, and conclusions or recommendations expressed in this material are those of the author(s) and do not necessarily reflect the views of the National Science Foundation. J.S.W. acknowledges support from the Wharton Dean’s Fund. J.M. and H.D. acknowledge support from R21 HD085201. A.R. and E.T. acknowledge support from NIH New Innovator Award DP2 OD008514, NIH/NCI PSOC award number U54 CA193417, NSF CAREER 1350601, NIH R33 EB019767, P30 CA016520, NIH 4DN U01 HL129998, NIH Center for Photogenomics RM1 HG007743, and the Charles E. Kauffman Foundation (KA2016-85223). S.S. acknowledges from NIH F30 AI114475. M.L acknowledges support from NIH R01GM108600, R01GM125301 and R01HL113147. R.B. acknowledges support from the NIH (DP2MH107055), the Searle Scholars Program (15-SSP-102), the March of Dimes Foundation (1-FY-15-344), a Linda Pechenik Montague Investigator Award, and the Charles E. Kauffman Foundation (KA2016-85223). This work used the Extreme Science and Engineering Discovery Environment (XSEDE), which is supported by National Science Foundation grant number OCI-1053575. Specifically, it used the Bridges system, which is supported by NSF award number ACI-1445606, at the Pittsburgh Supercomputing Center (PSC).

## Methods

### Data Pre-processing and Quality Control

SAVER can be applied to the matrix of raw UMI counts. However, in a standard scRNA-seq data set, many genes have zero total counts across all cells, or have non-zero count in at most 1 or 2 cells. Genes exhibiting such extremely sparse expression would not benefit from the SAVER procedure, since there is little data to form a good prediction; however these genes do not affect the estimates of the other genes, and thus are harmless if left in. As we show in Figure 9 of Supplementary Note 3, SAVER gives the most improvement for genes with medium to low expression, and for these extremely low abundance genes, the SAVER recovered values would be similar to the observed value. Thus, to reduce computational time, we recommend removing these genes at the start. There are several existing workflows^30–32^ that perform a conservative filtering of low abundance genes, which can be applied prior to application of SAVER.

### SAVER

Let *Y*_*gc*_ be the observed UMI count of gene *g* in cell *c*. We model *Y*_*gc*_ as a negative binomial random variable through the following Poisson-Gamma mixture

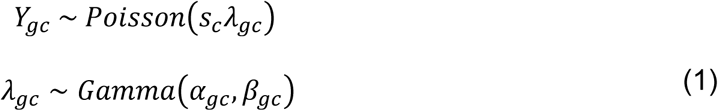

where *λ*_*gc*_ represents the normalized true expression. The Poisson model has been shown to be a good approximation of the noise in scRNA-seq data using UMIs^33,34^. Datasets without UMIs are subject to strong amplification bias and would violate the Poisson model assumed here. A gamma prior is placed on *λ*_*gc*_ to account for our uncertainty about its value. The shape parameter *α*_*gc*_ and the rate parameter *β*_*gc*_ are reparameterizations of the mean **μ**_*gc*_ and the variance *v*_*gc*_, see details in Supplementary Note 1. *s*_*c*_ represents the size normalization factor. In the following analyses, we use a library size normalization defined as the library size divided by the mean library size across cells, although other size factors such as those calculated by methods such as scran^12^, BASiCS^13^, SCnorm^35^, or through ERCC spike-ins can be used. SAVER can also accommodate pre-normalized data.

Our goal is to derive the posterior gamma distribution for *λ*_*gc*_ given the observed counts *Y*_*gc*_ and use the posterior mean as the normalized SAVER estimate 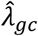. The variance in the posterior distribution can be thought of as a measure of uncertainty in the SAVER estimate. We adopt an empirical Bayes-like technique to estimate the prior mean and prior variance. First, we estimate the prior mean **μ**_*gc*_. We let **μ**_*gc*_ be a prediction for gene *g*derived from the expression of other genes in the same cell. Specifically, we use the log normalized counts of all other genes *g*′ as predictors in a Poisson generalized linear regression model with a log link function,

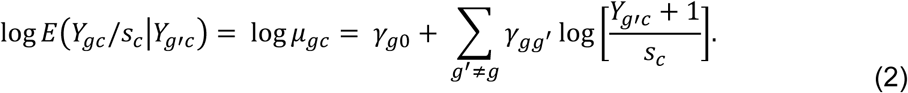

Since the number of genes often far exceeds the number of cells, a penalized Poisson Lasso regression is used to shrink most of the regression coefficients to zero. In a Lasso regression, a penalty parameter lambda is added to the likelihood to control the number of predictors that have nonzero coefficients. A large penalty would correspond to a model with very few nonzero coefficients while a small penalty would correspond to a model with many nonzero coefficients. The genes that have nonzero coefficients can be thought of as genes that are good predictors of the gene that is being estimated. We believe that this accurately reflects true biology since genes often only interact with a limited set of genes.

The regression is fit using the *glmnet* R package version 2.0-5^11^. For gene *g*, the response is the normalized observed expression 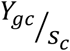 and the predictors are log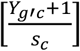 The regression model at the penalty with the lowest five-fold cross-validation error is selected (Supplementary Fig. 9). We then use the selected model to get our regression predictions 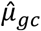, which we treat as the prior mean for each gene in each cell.

The next step is to estimate the prior variance by assuming a constant noise model across cells denoted by a dispersion parameter *ϕ*_*g*_ We consider three models for *ϕ*_*g*_: constant coefficient of variation 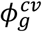 constant Fano factor 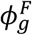 or constant variance 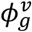. A constant coefficient of variation corresponds to a constant shape parameter *α*_*gc*_ = *α*_*g*_ in the gamma prior and a constant Fano factor corresponds to a constant rate parameter *β*_*gc*_ = *β*_*g*_ (see Supplementary Note 1). To determine which model for *ϕ*_*g*_ is the most appropriate, we calculate the marginal likelihood across cells under each model and select the one with the highest maximum likelihood, and then set 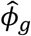 to the maximum likelihood estimate. Given 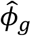 and the choice of noise model, we can derive 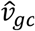

Now that we have both 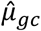 and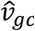, we can reparametrize, based on the chosen model for *ϕ*_*g*_, into the usual shape and rate parameters of the gamma distribution, 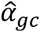 and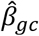. The posterior distribution is then

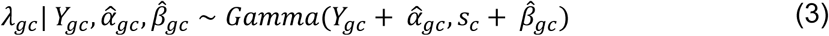

The SAVER estimate 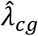 is the posterior mean, a weighted combination of the regression prediction and the normalized observed expression:

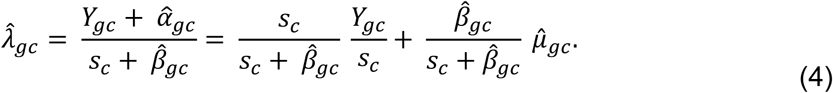

As seen from the above equation, the recovered expression 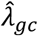 is a weighted average of the normalized observed counts *Y*_*gc*_ */s*_*c*_ and the prediction 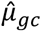. The weights are a function of the size factor *s*_*c*_ and, through the 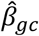 term, the gene’s predictability 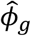 and its prediction *μ*_*gc*_. Genes for which the prediction is more trustworthy (small 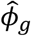) have larger weight on the prediction 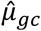. Genes with higher expression have larger weight on the observed counts and rely less on the prediction. Cells with higher coverage have more reliable observed counts and also rely less on the prediction. Supplementary Figure 10 shows example scenarios.

Estimating *ϕ*_*g*_ and computing the posterior distribution is fast computationally. The melanoma Drop-seq data with 12,241 genes and 8,498 cells took under 10 minutes total on one core of a standard desktop with an i7-3770 CPU. However, performing the prediction with the Lasso regression is computationally intensive. For the melanoma data, the Lasso regression took on average about 20 seconds per gene. However, this prediction step is highly parallelizable in the SAVER software and gene selection filters can be applied to reduce the dimensionality of the problem.

### Calculating correlations with SAVER

The SAVER estimate 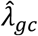 cannot be directly used to calculate gene-to-gene or cell-to-cell correlations since we need to account for its posterior uncertainty. Let the correlation between gene *g*and gene *g*′ be represented by *ρ*_*gg*′_ = *Cor*(***λ**_g_*, ***λ**_g′_*), where λ_*g*_ and λ_*g′*_ are the true expression vectors across cells. We can estimate *ρ*_*gg*′_ by calculating the sample correlation of the SAVER estimate 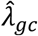 and scaling by an adjustment factor, which takes into account the uncertainty of the estimate:

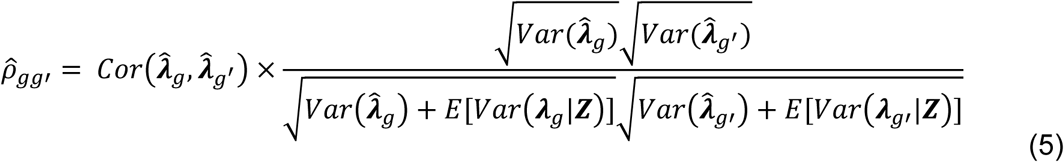

where *Var*(***λ***_*g*_ *|****Z***) is a vector of posterior variances. The same adjustment can be applied to cell-to-cell correlations. See Supplementary Note 2 for derivation of this adjustment factor.

### Distribution recovery

SAVER can be used to recover the distribution of either the absolute molecules counts or the relative expression values. Recovery of the absolute counts requires knowledge of the efficiency loss through ERCC spike-ins or some other control. To recover the absolute counts, we sample each cell from a Poisson-Gamma mixture distribution (i.e. negative binomial), where the gamma is the SAVER posterior distribution scaled by the efficiency. If the efficiency is not known or if relative expression is desired, we sample the expression level for each gene in each cell from the gene’s posterior gamma distribution.

### RNA FISH and Drop-seq analysis

The raw Drop-seq dataset contained 32,287 genes and 8,640 cells. Genes with mean expression less than 0.1 as well as cells with library size less than 500 or greater than 20,000 were removed. The filtered dataset contained 12,241 genes and 8,498 cells. RNA FISH measurements of 26 drug resistance markers and housekeeping genes were obtained across 7,000 to 88,000 cells from the same cell line. SAVER, MAGIC, and scImpute were performed on the Drop-seq data. MAGIC was performed using the Matlab version with default settings and library size normalization. scImpute version 0.0.2 was used with default settings. The 16 genes that were left after filtering are: 9 housekeeping genes (*BABAM1, GAPDH, LMNA, CCNA2, KDM5A, KDM5B, MITF, SOX10, VGF*) and 7 drug-resistance markers (*C1S, FGFR1, FOSL1, JUN, RUNX2, TXNRD1, VCL*).

Since the FISH and Drop-seq experiments have different technical biases, we normalized by a *GAPDH* factor for each cell, defined as the expression of *GAPDH* divided by the mean of *GAPDH* across cells in each experiment. *GAPDH* read counts have been used as a proxy for cell size^36^. Since some cells have very low or very high *GAPDH* counts, we filtered out cells in the bottom and top 10^th^ percentile. For the Gini coefficient analysis where we assume we do not know the efficiency, we sampled the SAVER dataset from the SAVER posterior gamma distributions. We then filtered out cells in the bottom and top 10^th^ percentile of *GAPDH* expression in the sampled SAVER dataset and normalized the remaining by the *GAPDH* factor. For the distribution recovery, we calculated the efficiency loss for each gene in each dataset as the mean FISH expression divided by the mean dataset expression. We scaled the Drop-seq, MAGIC, and scImpute dataset by the efficiency loss, filtered by *GAPDH*, and then normalized by the *GAPDH* factor. We scaled the SAVER posterior distributions by the efficiency loss and sampled from the Poisson-Gamma mixture to get the absolute counts as described above. We then performed the filtering and normalization by the *GAPDH* factor on the sampled SAVER dataset.

Correlation analysis was performed for pairs of genes in unnormalized FISH, Drop-seq, SAVER. Since the SAVER and MAGIC estimates were returned as library size normalized values, we rescaled by the library size to get the unnormalized values and used those to calculate the adjusted gene-to-gene correlations described above.

### Baron Study

Human pancreatic islet data contained 20,125 genes and 1,937 cells. Genes with mean expression less than 0.001 and non-zero expression in less than 3 cells were filtered out. The filtered dataset contained 14,729 genes and 1,937 cells. To generate the reference dataset, we selected genes that had non-zero expression in 25% of the cells and cells with a library size of greater than 5,000. We ended up with 2,284 genes and 1,076 cells.

### Chen Study

Mouse hypothalamus data contained 23,284 genes and 14,437 cells. Cells with library size greater than 15,000 were filtered out. Genes with mean expression less than 0.0002 and non-zero expression in less than 5 cells were filtered out. The filtered dataset contained 17,053 genes and 14,216 cells. To generate the reference dataset, we selected genes that had non-zero expression in 20% of the cells and cells with a library size of greater than 2,000. We ended up with 2,159 genes and 7,712 cells.

### La Manno Study

Human ventral midbrain data contained 19,531 genes and 1,977 cells. Genes with mean expression less than 0.001 and non-zero expression in less than 3 cells were filtered out. The filtered dataset contained 19,518 genes and 1,977 cells. To generate the reference dataset, we selected genes that had non-zero expression in 30% of the cells and cells with a library size of greater than 5,000. We ended up with 2,059 genes and 947 cells.

### Zeisel Study

Mouse cortex and hippocampus data contained 19,972 genes and 3,005 cells. To generate the reference dataset, we selected genes that had non-zero expression in 40% of the cells and cells with a library size of greater than 10,000 UMIs. We ended up with 3,529 genes and 1,800 cells. We also filtered out one cell that had abnormally low library size after gene selection to end up with 1,799 cells.

### Down-sampling datasets

Using the reference dataset as the true transcript count *λ*_*gc*_, we generated down-sampled observed datasets by drawing from a Poisson distribution with mean parameter *τ*_*c*_ *λ*_*gc*_, where *τ*_*c*_ is the cell-specific efficiency loss. To mimic variation in efficiency across cells, we sampled *τ*_*c*_ as follows,

1. 10% efficiency: *τ*_*c*_ ∼ *Gamma*(10, 100)
2. 5% efficiency: *τ*_*c*_ ∼ *Gamma*(10, 200)

The Baron, Chen, and La Manno datasets were sampled at 10% efficiency and the Zeisel dataset was sampled at 5% efficiency.

### Implementation of methods on down-sampled data

We compared the performance of SAVER against using the library-size normalized observed dataset, MAGIC, and scImpute. The missing data imputation techniques were performed on the library size normalized observed data treating zeros as missing. KNN imputation was performed using the *impute.knn* function in the *impute* R package version 1.48.0, with parameters *rowmax* = 1, *colmax* = 1, and *maxp = p*. SVD imputation was performed on the row and column centered matrix using the *soft.Impute* function in the *softImpute* R package version 1.4, with parameters *rank.max =* 50, *lambda =* 30, and *type =* “svd”. Random forest imputation was performed on the matrix transpose with the *missForest* R package version 1.4 with default parameters.

Percentage change over observed was defined as

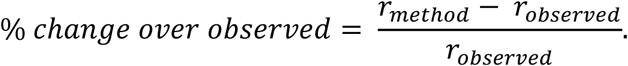

### Gene-to-gene and cell-to-cell correlation analysis

Pairwise Pearson correlations were calculated for each library size normalized dataset and imputed dataset. Since the SAVER estimates have uncertainty, we want to calculate the correlation based on *λ*_*gc*_. Correlations were first calculated using the SAVER recovered estimates 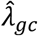 and scaled by the correlation adjustment factor described above.

The correlation matrix distance (CMD) is a measure of the distance between two correlation matrices with range from 0 (equal) to 1 (maximum difference)^24^. The CMD for two correlation matrices ***R***_1_, ***R***_2_ is defined as

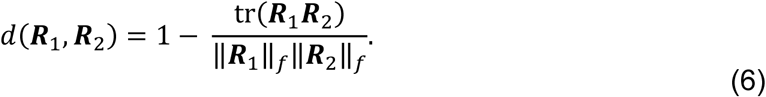

### Differential expression analysis of down-sampled datasets

For each down-sampled dataset, ten SAVER sampled datasets were generated by sampling from the posterior gamma distribution. A Wilcoxon rank sum test was run on each of the sampled datasets and the combined p-value was obtained via Rubin’s rules for multiple imputation^37^. FDR control was set to 0.01 and no fold change cutoff was used. MAST version 1.0.5 was run on the library size normalized expression counts with the condition and scaled cellular detection rate as the Hurdle model input. The combined Hurdle test results were used. scDD version 1.2.0 was run on the library size normalized expression counts with default settings. Both the nonzero and the zero test results were used. SCDE version 2.2.0 was run on unnormalized expression counts with default parameters, except number of randomizations was set to 100. The p-value was calculated according to a two-sided test on the corrected Z-score.

To calculate the estimated false discovery rate, we first performed a permutation of the cell labels and determined the number of genes called as differentially expressed according to the p-value threshold defined for the unpermuted data. This number divided by the number of differentially expressed genes in the unpermutated data is the false discovery rate for that one permutation. The final estimated false discovery rate is the average of the false discovery rates over 20 permutations. For SAVER, one sampled dataset was considered one permutation.

### Cell clustering and t-SNE visualization

Seurat version 2.0 was used to perform cell clustering and t-SNE visualization following the workflow detailed at http://satijalab.org/seurat/pbmc3k_tutorial.html. Briefly, normalization without filtering, identification of highly variable genes, scaling, PCA, jackStraw, cell clustering, and t-SNE were applied to the reference, down-sampled, SAVER, MAGIC, and scImpute datasets. The number of principal components used for cell clustering and t-SNE were identified through the jackStraw procedure. For the reference datasets, 15 PCs were chosen for Baron, Chen, and La Manno and 20 PCs were chosen for Zeisel. The number of principal components chosen for each down-sampled dataset and method is shown in Supplementary Figure 8. The resolution for each reference dataset was chosen such that the cell clustering had the most agreement with the t-SNE visualization. Resolutions of 0.7, 0.6, 1.1, and 0.8 were chosen for Baron, Chen, La Manno, and Zeisel reference datasets respectively. Cell clusterings were calculated for each observed and recovered dataset at resolutions of 0.4-1.4 at intervals of 0.1. The Jaccard index was calculated at each resolution with the reference dataset, and the maximum Jaccard index was then reported. The Jaccard index was calculated using the R package *clusteval* version 0.1.

### Hrvatin Study

Mouse visual cortex data contained 25,187 genes and 65,539 cells. Genes with mean expression less than 0.00003 and non-zero expression in less than 4 cells were filtered out. The filtered dataset contained 19,155 genes and 65,539 cells. 47,209 cells were classified into cell types by the authors. SAVER was run on a subsample of 10,000 cells. Out of these 10,000 cells, 7,387 cells had a subtype label and Seurat was used to cluster these cells. 35 principal components were chosen for the observed data and 30 principal components were chosen for the SAVER results as determined by the jackstraw procedure.

